# RPM: an open-source rotation platform for open- and closed-loop vestibular stimulation in head-fixed mice

**DOI:** 10.1101/2023.05.19.541416

**Authors:** Xavier Cano-Ferrer, Alexandra Tran-Van-Minh, Ede Rancz

## Abstract

Head fixation allows the recording and presentation of controlled stimuli and is used to study neural processes underlying spatial navigation. However, it disrupts the head direction system because of the lack of vestibular stimulation.

To overcome this limitation, we developed a novel rotation platform which can be driven by the experimenter (open-loop) or by animal movement (closed-loop). The platform is modular, affordable, easy to build and open source. Additional modules presented here include cameras for monitoring eye movements, visual virtual reality and a micro-manipulator for positioning various probes for recording or optical interference.

We demonstrate the utility of the platform by recording eye movements and showing the robust activation of head-direction cells. This novel experimental apparatus combines the advantages of head fixation and intact vestibular activity in the horizontal plane. The open-loop mode can be used to study e.g. vestibular sensory representation and processing, while the closed-loop mode allows animals to navigate in rotational space, providing a better substrate for 2-D navigation in virtual environments. Documentation is available at (https://ranczlab.github.io/RPM/).

## 1 Introduction

Head-fixation of mice proved exceedingly useful in recent years to allow acute electrophysiological and optical recordings in awake, behaving animals [1, 2]. Furthermore, it facilitates camera-based tracking of behaviourally relevant features (e.g. eye movements) and allows for the controlled delivery of sensory stimuli and rewards. On the other hand, head-fixation disables the vestibular system, a key sensory component underlying navigation [3], largely limiting experimental environments to one-dimensional corridors [4]. Activation of the head-direction (HD) network is particularly needed for spatial navigation [5, 6] and depends on information from the horizontal semicircular canals [7]. Thus the lack of vestibular input evoked by horizontal rotation is a plausible reason for a lower percentage of place cells in head-fixed, virtual environments [8, 9]. While using a floating real-world environment improves spatial information rates in hippocampal place cells [10] compared to visual VR environments, the head direction system remains impaired due to the lack of vestibular input. Our motivation for developing a novel rotation platform was to control horizontal angular vestibular input in head-fixed mice. Previous approaches were limited to passive rotation [11] and thus lacked the ability of the animals to control rotation, i.e. navigate. When navigation in rotation space was allowed by fixing the animal’s head to a bearing [12], there was no option to control the rotation either passively or actively. An elegant alternative approach was used by Voigts et al. [13] where headpost torque was used to read out the animal’s intention to rotate its head and fed back through a rotational motor. However, this approach is difficult to combine with visual virtual reality, requires frequent calibration, is not documented in detail and requires significantly more expertise to build. Here we present an open-source rotational platform which can be used to control and interfere with the sensory environment. We provide documentation for building it and two use cases for studying vestibular evoked eye movements and head-direction cell activity during head-fixed, animal-controlled rotation.

## 2 Materials and Methods

### 2.1 Animals and headplate surgery

All animal experiments were prospectively approved by the local ethics panel of the Francis Crick Institute (previously National Institute for Medical Research) and the UK Home Office under the Animals (Scientific Procedures) Act 1986 (PPL: 70/8935). All surgery was performed under isoflurane anaesthesia, and every effort was made to minimize suffering. Animals were housed in individually ventilated cages under a 12 hr light/dark cycle. Briefly, the scalp was opened with an incision in the midline and connective tissue covering the bone was removed. An aluminium head plate was secured to the dorsal surface of the skull using dental acrylic, and the skin was fixed to it using VetBond. Animals were allowed to recover for 5 days before any experiments.

### 2.2 Eyemovement recordings

A virtual drum consisting of gratings with a frequency of 0.16 cycles per degree (cpd) was displayed on 4 monitors (Dell U2412B) around the setup and drifted during the entire recording at 1Hz. The vestibular stimulus was delivered by the rotation platform in clockwise (CW) and counterclockwise (CCW) directions at 40 °/s. Images were recorded using an ELP-USBFHD01M web camera at 30 Hz. The pupil outlines were detected as illustrated in Figure 4b using DeepLabCut [14]. Briefly, 8 points were detected on the edge of the pupil (red circles), onto which an ellipse was fitted (cyan line). The pupil position was defined as the centre of the ellipse (red square). Data were acquired and analysed using custom-written Python scripts.

### 2.3 HD cell recordings

We followed the recording method from [15]. We made post-subicular recordings (AP: -4.25 mm; ML: -1 to -2.5 mm; DV 1.25 to 1.75 mm from pia) from 2 adult male mice for at least five full bidirectional rotations in passive and at least 1 (but typically 3-4) full cycles in active rotation. Extracellular recordings were made using a tetrode-arranged silicone probe (A8×1-tet-2mm-200-121, Neuronexus), connected to a digitizing headstage (RHD2132, Intan technologies) via an adaptor (Adpt.A32-Omnetics32, Neuronexus). Voltage signals were filtered (0.1 – 7.5 kHz) and digitized at 24414 Hz using an LR10 interface and SynapseLite software (Tucker-Davies Technologies, TDT). Data were extracted using the TDT python SDK, spikes were detected and clustered using klusta [16] and further analysed using custom-written Python scripts. Tuning curves were calculated at 1° resolution and smoothed (6°, Gaussian). HD cells were defined as in [17, 15]. Briefly, a von Mises function was fitted to the tuning curve and only units with *κ* > 1, peak firing rates > 1 Hz and probability of non-uniform distribution < 0.001 (Rayleigh test) were included.

## 3 Results

### 3.1 Design criteria

We set out to develop a rotation platform capable of open-loop and closed-loop vestibular stimulation for mice which are head-fixed but able to move on an air-supported spherical treadmill. Principal requirements were: 1. accurate tracking of the movement of mice; 2. control of the platform rotation in open- and closed-loop modes, and 3. compatibility with physiological and behavioural recording modalities. Furthermore, we aimed to develop a modular, low-cost and open-source device.

### 3.2 Implementation

We provide full documentation, including assembly instructions, design choices and characterization of the components and the platform, as well as software requirements and code at https://ranczlab.github.io/RPM/.

In brief, the movement of the spherical treadmill is read out by optical sensors and fed through the rotary joint to connect to a microcontroller (PJRC Teensy^®^ 3.5). In addition, the microcontroller receives information from a homing sensor and a rotary encoder (Figure 1a). The microcontroller, in turn, controls the platform movement through the motor controller (Figure 1b), and outputs the position of the spherical treadmill and the platform for recording. Other signals recorded on the platform, e.g. from cameras, or signals to the platform to control additional devices, can also be fed through the rotary joint (Figure 1c).

**Figure 1:**
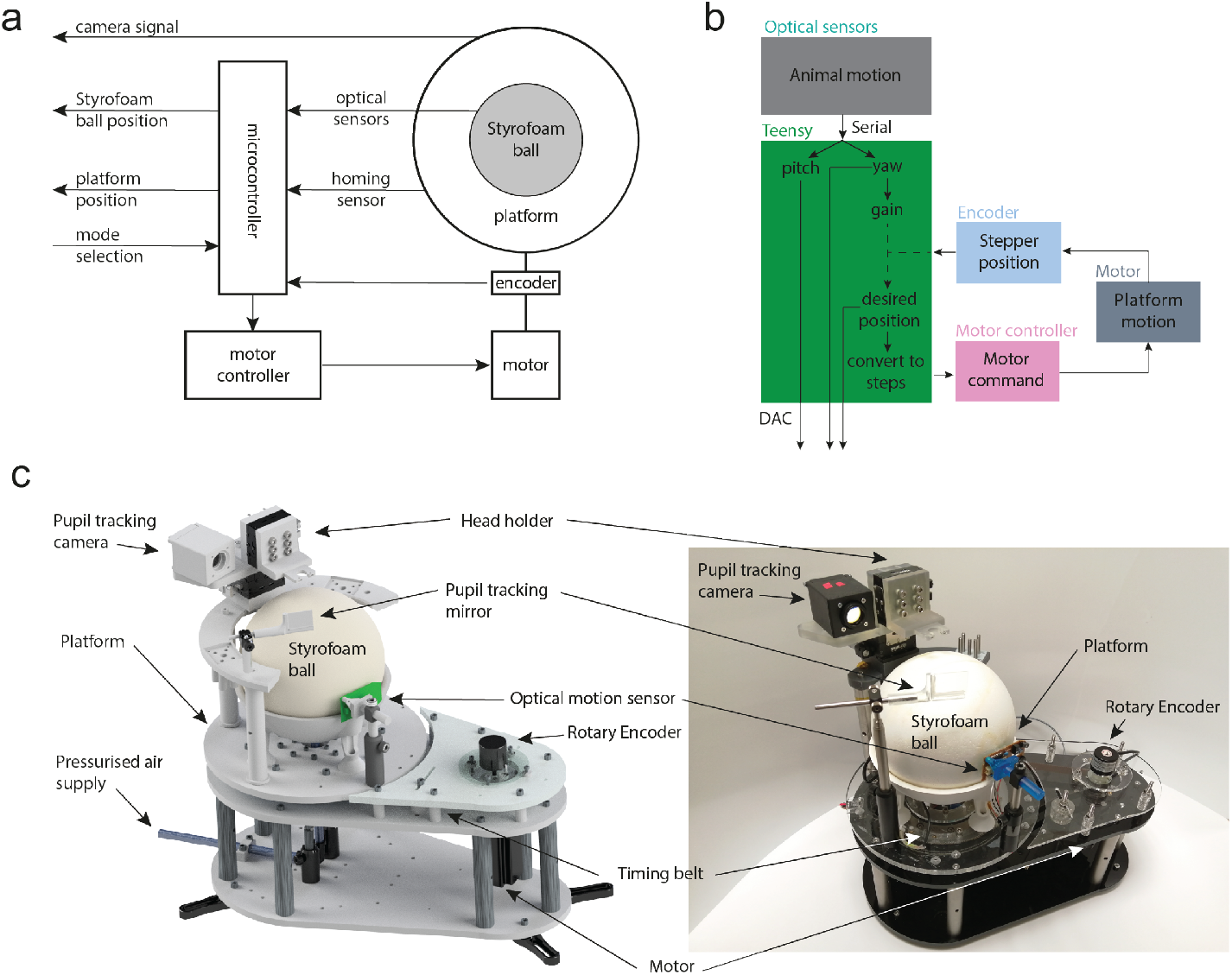
RPM: Rotation Platform for Mice. (A) High-level I/O, (B) algorithmic implementation and (C) RPM device, 3D model (left) and photograph (right).

### 3.3 Characterization and validation of platform in closed and open loop

#### 3.3.1 No load performance, minimal incremental motion and back-lash

The rotary platform’s accuracy and repeatability have been evaluated by recording the platform’s actual position using an incremental encoder and comparing it to the target position set by the microcontroller. Figure 2a shows an example test of the stage performance without the load of the head-plate holder. The stage is repeatedly sent to two set points (0 and 360 degrees). The final positions are highly accurate (360.06 ± 0.11° and 0.04 ± 0.07°), suggesting little error accumulation. This data also shows that the stepper counter in the microcontroller provides an accurate measure of the platform position. Further characterization was carried out with a typical load condition (i.e. head-plate holder included).

**Figure 2:**
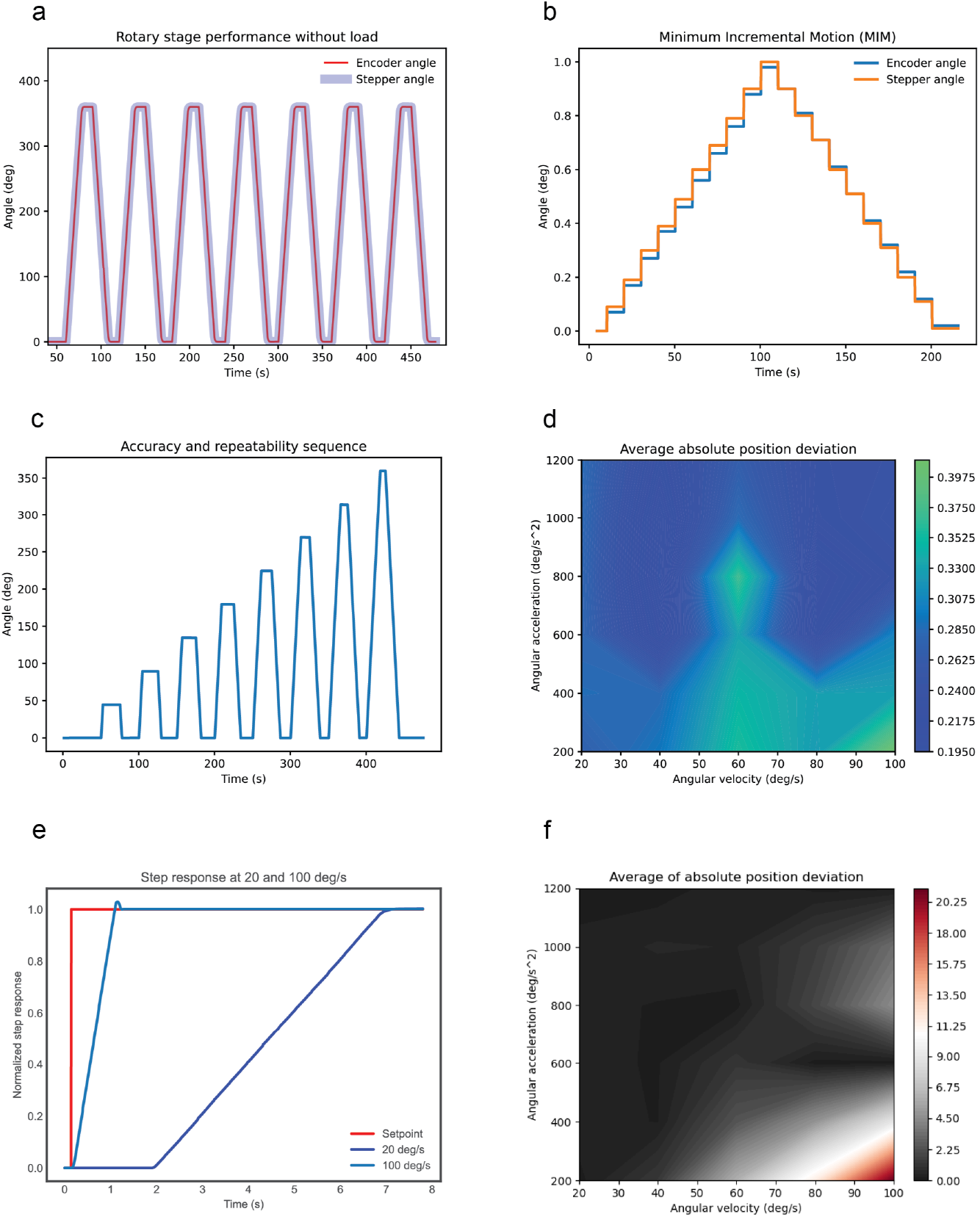
Rotary stage performance. (A) Rotary encoder and stepper motor signals during full revolution cycles. (B) Minimum incremental motion measurement. (C) A sequence of bi-directional movements to measure accuracy and repeatability. (D) Average absolute position deviation across all angular speed and acceleration conditions tested. (E) Example step responses at two different angular velocities (20 and 100 °·s^-1^), (F) Position deviation map across all angular velocity/acceleration settings.

The measured minimum incremental motion (MIM) was 0.1° (Figure 2b). The backlash, in the worst-case scenario corresponding to the offset created by the change in the direction of the MIM tests, has been 0.04°.

#### 3.3.2 Accuracy and repeatability

In order to evaluate the accuracy and repeatability of the rotary platform, a sequence of increments of 45°. in both rotation directions have been performed, collecting 8 position deviations in each direction in a trapezoidal motion profile (Figure 2c). The results show a high directional accuracy (0.37°) and repeatability (0.10°) in experimental load conditions. Figure 2d show the distribution of the absolute average position deviation mapped across the range of angular velocity and acceleration values (20-100 °·s^-1^ and 200-1200 °·s^-2^). The equations for the calculation of these measures can be found in the methods section.

#### 3.3.3 Closed-loop stability

The closed-loop algorithm drives motor rotation in response to the motion sensed by the optical sensor, i.e. due to the Styrofoam ball rotation actuated by the animal. The system thus has to respond to rapid changes in the target position. We have simulated sudden and unpredictable changes in the input angle as a step function of 90°. The experiment has been performed with the same load conditions as the closed-loop algorithm. Stability has been evaluated by the analysis of the step response for all previously described conditions (20-100 °·s^-1^ and 200-1200 °·s^-2^, Figure 2e). The contour plot has been extracted to be used as a map to avoid regions of inaccuracy and instability. Sometimes even achieving a small error, the stage can overshoot, as shown in Figure 2e. The regions with more instability correspond to the regions where the average absolute position deviation is higher (Figure 2f). This behaviour usually corresponds to conditions of high speed and not high enough acceleration. These performance measures should be conducted whenever the load or load distribution on the platform is altered.

**Table 1.** Rotary stage specifications.

|  | Value | Units |
| --- | --- | --- |
| Motor resolution | 0.0024 | ° |
| Encoder resolution | 0.025 | ° |
| Accuracy | 0.37 | ° |
| Repeatability | 0.10 | ° |
| Minimum incremental motion | 0.10 | ° |
| Backlash | 0.04 | ° |
| Maximum angular velocity tested | 100 | °·s<sup>-1</sup> |
| Maximum angular acceleration tested | 1200 | °·s<sup>-2</sup> |

### 3.4 Add-ons

A ring-shaped instrument platform can be mounted around the sphere to hold several devices, such as an animal head-holder, cameras, and manipulators for electrical recording and optogenetic stimulation. Below we present options for head-plate holders and instruments for manipulating recording probes or optical fibres, as well as for installing cameras and video screens for eye-movement tracking and virtual navigation, respectively.

#### 3.4.1 Head-holder

We designed a head fixation system with three degrees of freedom along three perpendicular translational axes (Figure 3a). The height of the head-plate holder can be adjusted to accommodate mice of different sizes. In addition, two axes can be used to adjust the position of the animal’s head in the horizontal plane. This is particularly useful in vestibular experiments to centre the platform rotation axis on the vestibular organ of interest. The head-holding system consists of 3 manual translational stages (Thorlabs Inc. MT1/M) to ensure reliability and repeatability across experiments. The resolution of the system is 10 microns with a range of 13 mm in all directions. The stages are connected using a 3D printed linker, and various head-plate holders can be attached to the vertical stage (e.g. for attaching micro-manipulators and cameras).

**Figure 3:**
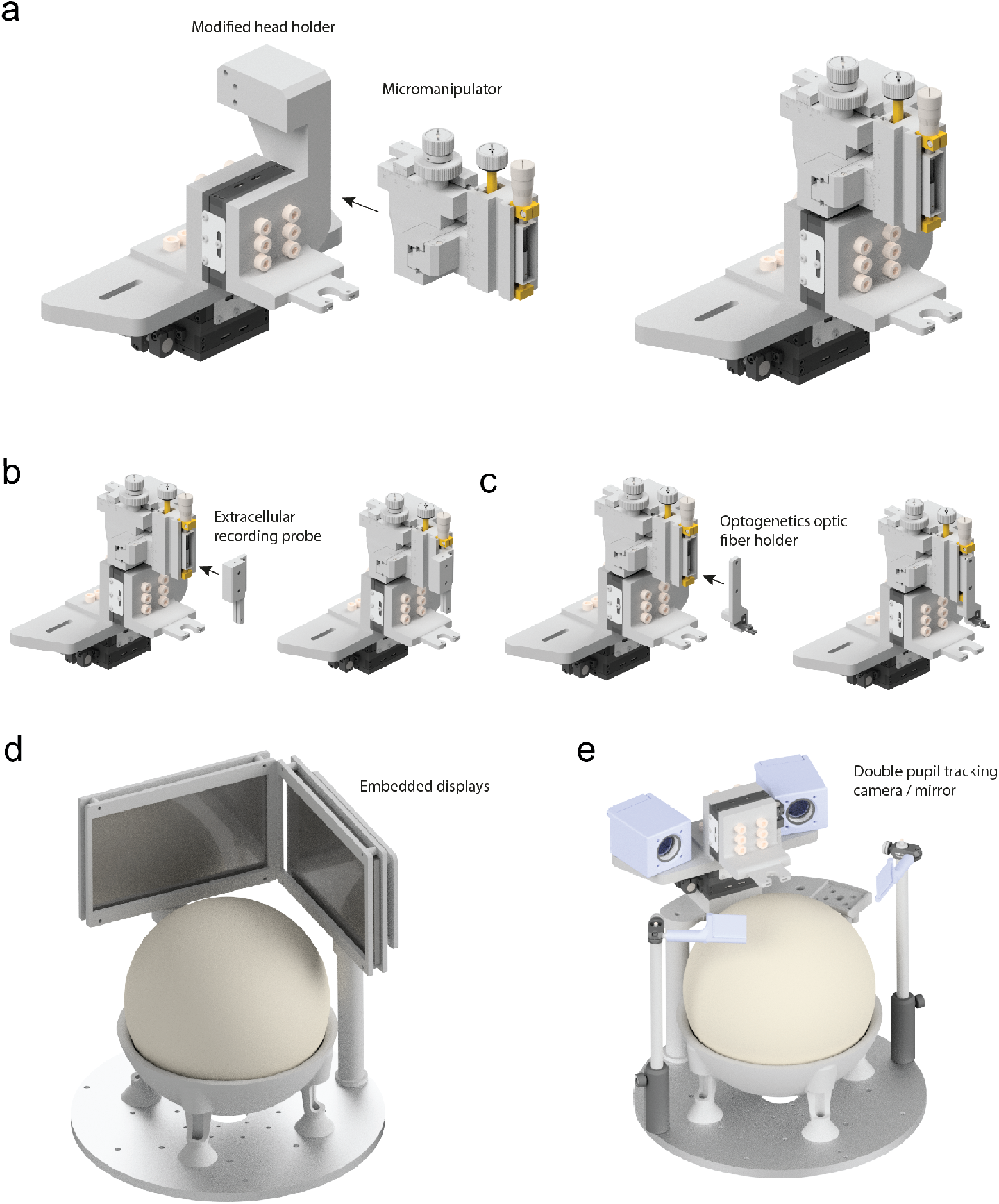
Platform add-ons. (A) 3D printable head-plate holder modified to attach a micro-manipulator, (B) a micro-manipulator with the extracellular recording probe holder, (C) a micro-manipulator in the optogenetics fibre holder configuration, (D) the rotary platform with two embedded displays, (E) head-plate holder modified to have two pupil tracking cameras and the rotary platform with two hot mirrors.

#### 3.4.2 Micro-manipulator and probe holders

The head-holder in Figure 3a was designed to attach a manual micro-manipulator (Märzhäuser Wetzlar MM33) and provide a capability for attaching e.g. electrophysiological (Figure 3b) or optical (Figure 3c) probe holders. The device provides fine adjustment (0.1 mm) on the three axes with a finer adjustment (0.01 mm) for the vertical axis and a travel range of 37,20 and 25 mm for the x,y, and z axes, ideal for precise adjustment of the position and depth of various probes.

#### 3.4.3 Cameras

On-platform cameras provide useful capability to image e.g. eye-, face- or body movements. We have implemented a camera system to monitor pupil movements (Figure 3e). The animal’s pupil is illuminated by an infrared LED, and the image is reflected from a hot mirror (Thorlabs Inc. FM02R) with an incidence angle of 45°. Both the mirror and camera position and angle can be adjusted for adequate positioning. An IR-sensitive camera with USB connectivity (ELP-USBFHD01M) was placed behind a long-pass filter (Thorlabs Inc. FEL0750) to record the image of the pupil without contamination from other light sources.

#### 3.4.4 Embedded displays and virtual reality

We attached two 8” TFT displays to the rotary platform using Thorlabs RS6P/M posts, 3D printed parts and a laser-cut display frame (Figure 3d). The displays can be used to display visual stimuli and allow navigation in a 2D virtual environment.

The VR environment runs in Godot on a Raspberry Pi 4 Model B running Raspbian 10 (Buster). The single board computer was placed on the platform and connected to the screens via micro HDMI cables, while power was provided through the rotary joint. The optical sensor inputs transformed on the microcontroller to calculate the motor displacement were converted into HID commands on the PCB and transmitted to the Raspberry Pi via one of the USB connections of the rotary ring. The environment consisted of a virtual corridor, with regularly placed white cues on the right-hand side wall and terminating with a uniform green wall. The displacement of the ball measured by the optical sensors controlled both the angular displacement of the platform and the navigation within the virtual environment. An example video (Video 1) shows navigation in the virtual environment and the physical rotation of the platform. The trajectory of the virtual position of the animal is plotted in red in the bottom right corner.

### 3.5 Use cases

#### 3.5.1 Eye-movement recording during open-loop configuration

We carried out pupil recordings to illustrate the utility of the platform for vestibular and visually evoked eye movements. Visual stimuli were provided by 4 screens surrounding the platform, and the pupil position was tracked using an on-platform camera (Figure 4a). The position of the pupil was detected using a deep-learning-based approach[14] (Figure 4b). Drifting gratings were used to evoke the optokinetic reflex, while platform rotation was used to evoke the vestibulo-ocular reflex simultaneously. Eye movements showed typical nystagmus beating in the direction of the physical rotation (Figure 4c-d). These data demonstrate the utility of the platform for open-loop vestibular stimulation and eye tracking.

**Figure 4:**
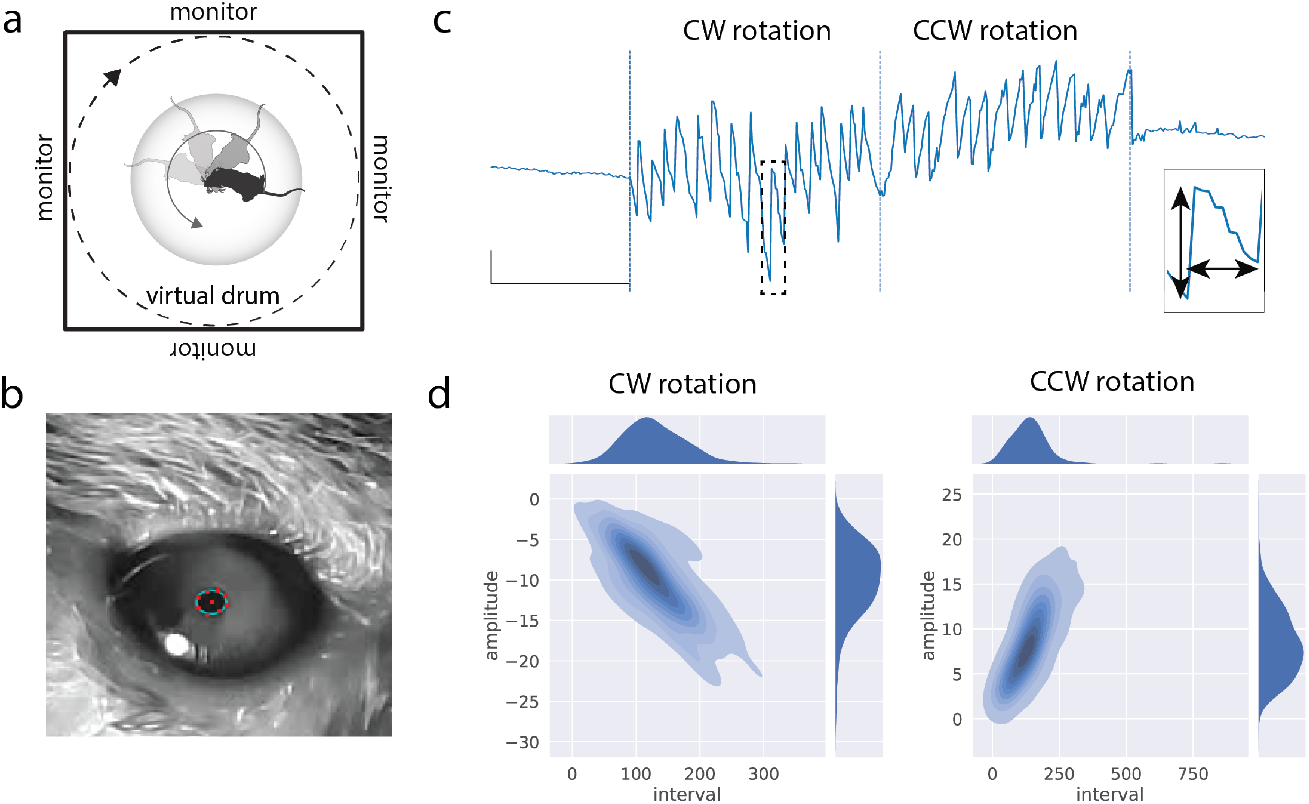
Analysis of vestibulo-visual evoked eye-movements. (A) Recording configuration. (B) Example camera image and detection points. (C) Example trace showing pupil movement (y-axis) during clockwise (CW) and counterclockwise (CCW) platform rotation. Scale bar: 4 pixels, 1s. Insert: amplitude and interval measures on a single saccade. (D) Distribution of peaks saccade amplitudes vs duration. Average of 16 trials across 4 different mice

#### 3.5.2 Extracellular recording in open- and closed-loop configuration

Next, we have set out to test the suitability of the vestibular platform for electrophysiological recordings. We recorded extracellular voltage signals with acutely inserted silicone probes. We targeted the postsubiculum (PoS) in awake mice during passive (open-loop, position controlled by experimenter) and active (closed-loop, mouse movement determines position) rotation. The postsubiculum was chosen as it has been reported to contain head-direction cells in large proportion [18]. In turn, HD cells are known to depend on intact vestibular activity [7] and can be activated by passive rotation as well [19]. The rotation platform was placed in the middle of a black cylinder containing one salient visual landmark. Animals were familiarised with the apparatus and the environment before the recordings. Example traces of unit (0.3 – 5 kHz) and LFP (1 – 100 Hz) recordings as well as position, velocity and acceleration, are shown in Figure 5a. There were no discernible artefacts or changes in noise level at the onset of motion or rapid change in direction. We could readily record (HD) cells (n = 7 cells from 2 animals) from the PoS. We identified HD cells using established criteria (see [15] and methods) during passive rotation with a range of ± 360° (Figure 5b). Next, we allowed animals to actively rotate at least one (but typically 3-4) full rotations. Directional tuning curves between active and passive rotation showed no significant difference in the preferred direction (Figure 5c-d; mean difference -18 ± 25°, n = 7, p = 0.1). Similarly, maximum firing rates between active and passive conditions were not significantly different (Figure 5e; mean firing rate 15.5 ± 5.5 Hz vs 15.7 ± 6.8 Hz, n = 7, p = 0.9). We can thus conclude that the vestibular platform is suitable for electrophysiological recordings, and HD activity is present during both active and passive rotation.

**Figure 5:**
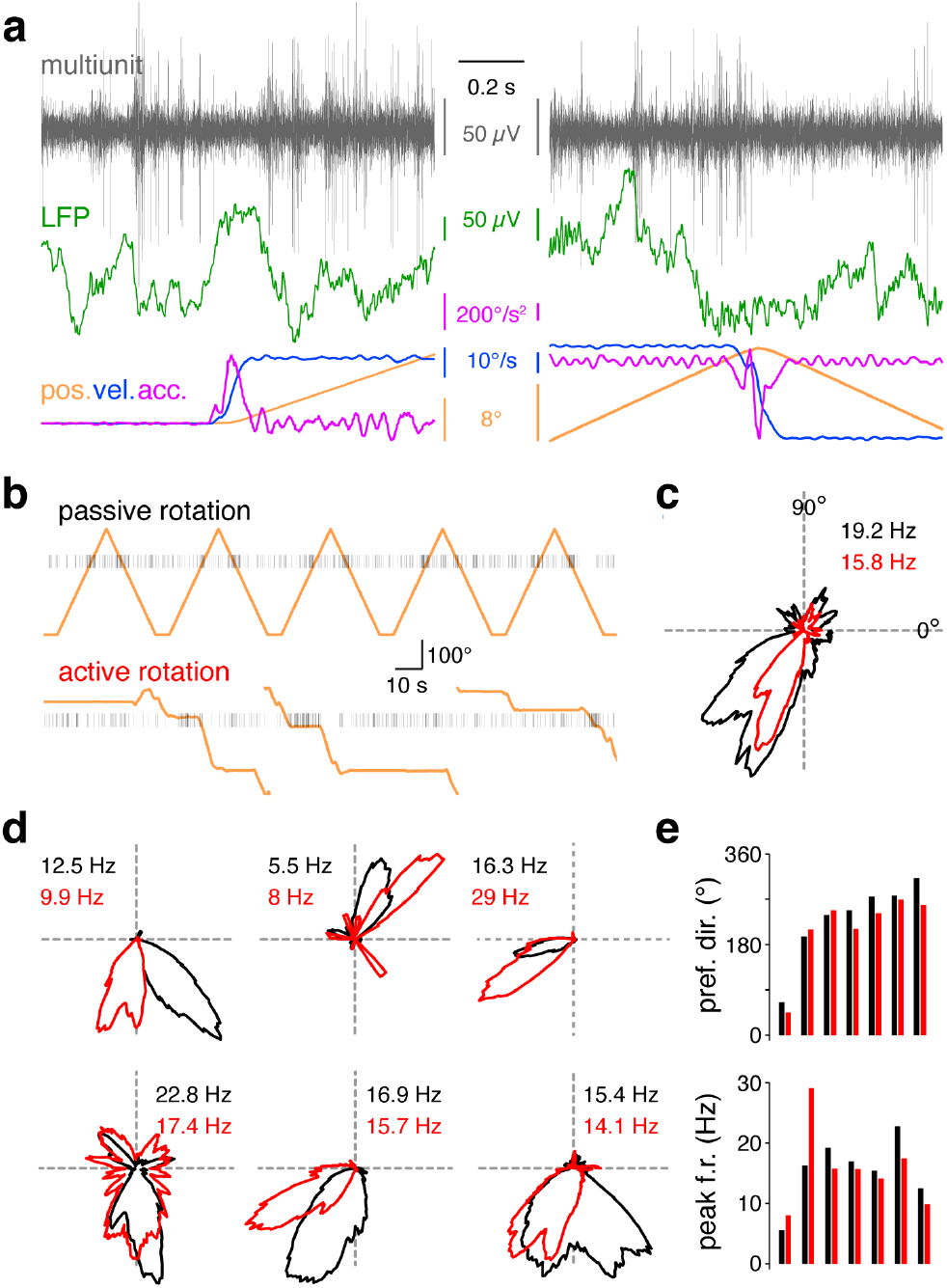
Extracellular recordings during rotation. A. Example extracellular voltage traces showing multiunit (0.3 – 5 kHz) and LFP (1-100 Hz) activity recorded during the start of movement (left) or an abrupt change of direction (right). Note the lack of discernible artefacts. B. Spike positions from an isolated single unit during passive and active rotation. C. Polar plot of firing rate as a function of direction for the unit in B. Peak firing rates are given. Black: passive rotation. Red: active rotation. D. Polar plots of firing rate for a further 6 HD tuned units. E. Bar graph showing preferred head direction (top) and peak firing rate (bottom) between passive (black) and active (red) rotation.

## 4 Discussion

There are inherent limitations in angular velocities and acceleration achievable by the rotation platform, depending e.g. on the weight and distribution of the platform assembly. Furthermore, animals of higher weight can also cause oscillation as their posterior part can swivel the ball during a change of direction. This can be counteracted by installing guard walls at the sides of the head holder assembly, keeping the mouse slightly constrained. Average and sustained head motion velocities in natural self-motion are relatively slow (±150 °·s^-1^, [20]); nevertheless, mice can experience up to 1300 °·s^-1^ angular velocity [21]. The rotation platform and assembly presented here can reliably reach up to 100 °·s^-1^ angular velocity and 1200 °·s^-2^ acceleration. This is adequate for most experiments, as published studies typically use up to 80 °·s^-1^ [22, 23]. Finally, We show that the HD network is robustly activated during self-guided rotation. It is important to note that the HD system also receives proprioceptive and motor efferent signals [24], which may be different on the device presented (especially regarding head-on-body motion).

In summary, we provide an open-source device with build documentation, which can serve as a starting point for researchers interested in performing both closed and open-loop vestibular experiments. We also provide various alternatives on how to adapt the experimental setup to achieve different experimental scenarios.

## 5 Supplementary Information

**Equation 1**. Accuracy

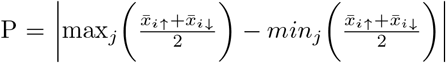

Where 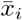 is the average inaccuracy at position i. Up and down arrows represent the positive and negative direction.

**Equation 2**. Repeatability

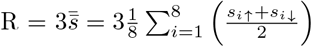

Where *s*_*i*_ is the standard deviation at position i. Up and down arrows represent the positive and negative direction. Repeatability is specified at a factor of 3 standard deviations (+/-1.5 sigma).

## 6 Acknowledgement

We thank Bogdan A. Horobet and Andrew Erskin for help with the development of the first prototype. We thank Renato Delaqua and Alan Ling from the Francis Crick Institute Mechanical Engineering Workshop for help in manufacturing some components. This work was supported by the Wellcome Trust (104285/B/14/Z) and the Francis Crick Institute, which receives its core funding from Cancer Research UK (FC001153), the United Kingdom Medical Research Council (FC001153), and the Wellcome Trust (FC001153). A.TVM. received funding from the European Union’s Horizon 2020 research and innovation programme under the Marie Sklodowska-Curie grant agreement No 747902. The funders had no role in study design, data collection and analysis, decision to publish, or preparation of the manuscript.

## References

[1] Lloyd E. Russell, Henry W. P. Dalgleish, Rebecca Nutbrown, Oliver M. Gauld, Dustin Herrmann, Mehmet Fişek, Adam M. Packer, and Michael Häusser. All-optical interrogation of neural circuits in behaving mice. Nature Protocols, 17(7):1579–1620, July 2022.

[2] Fritjof Helmchen, Ariel Gilad, and Jerry L. Chen. Neocortical Dynamics During Whisker-Based Sensory Discrimination in Head-Restrained Mice. Neuroscience, 368:57–69, January 2018.

[3] Paul F. Smith, Cynthia. L. Darlington, and Yiwen Zheng. Move it or lose it-Is stimulation of the vestibular system necessary for normal spatial memory? Hippocampus, pages NA–NA, 2009.

[4] Matthias Minderer, Christopher D. Harvey, Flavio Donato, and Edvard I. Moser. Virtual reality explored. Nature, 533(7603):324–325, May 2016.

[5] Brett Gibson, William N. Butler, and Jeffery S. Taube. The Head-Direction Signal Is Critical for Navigation Requiring a Cognitive Map but Not for Learning a Spatial Habit. Current Biology, 23(16):1536–1540, August 2013.

[6] S. S. Winter, B. J. Clark, and J. S. Taube. Disruption of the head direction cell network impairs the parahippocampal grid cell signal. Science, 347(6224):870–874, February 2015.

[7] Stephane Valerio and Jeffrey S. Taube. Head Direction Cell Activity Is Absent in Mice without the Horizontal Semicircular Canals. The Journal of Neuroscience: The Official Journal of the Society for Neuroscience, 36(3):741–754, January 2016.

[8] Zahra M Aghajan, Lavanya Acharya, Jason J Moore, Jesse D Cushman, Cliff Vuong, and Mayank R Mehta. Impaired spatial selectivity and intact phase precession in two-dimensional virtual reality. Nature Neuroscience, 18(1):121–128, January 2015.

[9] Guifen Chen, John A. King, Neil Burgess, and John O’Keefe. How vision and movement combine in the hippocampal place code. Proceedings of the National Academy of Sciences, 110(1):378–383, January 2013.

[10] Mary Ann Go, Jake Rogers, Giuseppe P. Gava, Catherine E. Davey, Seigfred Prado, Yu Liu, and Simon R. Schultz. Place Cells in Head-Fixed Mice Navigating a Floating Real-World Environment. Frontiers in Cellular Neuroscience, 15:618658, February 2021.

[11] Mateo Vélez-Fort, Charly V. Rousseau, Christian J. Niedworok, Ian R. Wickersham, Ede A. Rancz, Alexander P.Y. Brown, Molly Strom, and Troy W. Margrie. The Stimulus Selectivity and Connectivity of Layer Six Principal Cells Reveals Cortical Microcircuits Underlying Visual Processing. Neuron, 83(6):1431–1443, September 2014.

[12] Guifen Chen, John Andrew King, Yi Lu, Francesca Cacucci, and Neil Burgess. Spatial cell firing during virtual navigation of open arenas by head-restrained mice. eLife, 7:e34789, June 2018.

[13] Jakob Voigts and Mark T. Harnett. Somatic and Dendritic Encoding of Spatial Variables in Retrosplenial Cortex Differs during 2D Navigation. Neuron, 105(2):237–245.e4, January 2020.

[14] Tanmay Nath, Alexander Mathis, An Chi Chen, Amir Patel, Matthias Bethge, and Mackenzie Weygandt Mathis. Using DeepLabCut for 3D markerless pose estimation across species and behaviors. Nature Protocols, 14(7):2152–2176, July 2019.

[15] Adrien Peyrache, Natalie Schieferstein, and Gyorgy Buzsáki. Transformation of the head-direction signal into a spatial code. Nature Communications, 8(1):1752, December 2017.

[16] Cyrille Rossant, Shabnam N Kadir, Dan F M Goodman, John Schulman, Maximilian L D Hunter, Aman B Saleem, Andres Grosmark, Mariano Belluscio, George H Denfield, Alexander S Ecker, Andreas S Tolias, Samuel Solomon, György Buzsáki, Matteo Carandini, and Kenneth D Harris. Spike sorting for large, dense electrode arrays. Nature Neuroscience, 19(4):634– 641, April 2016.

[17] Adrien Peyrache, Marie M Lacroix, Peter C Petersen, and György Buzsáki. Internally organized mechanisms of the head direction sense. Nature Neuroscience, 18(4):569–575, April 2015.

[18] Kathleen E Cullen and Jeffrey S Taube. Our sense of direction: progress, controversies and challenges. Nature Neuroscience, 20(11):1465–1473, November 2017.

[19] Michael E. Shinder and Jeffrey S. Taube. Active and passive movement are encoded equally by head direction cells in the anterodorsal thalamus. Journal of Neurophysiology, 106(2):788–800, August 2011.

[20] Arne F. Meyer, Jasper Poort, John O’Keefe, Maneesh Sahani, and Jennifer F. Linden. A Head-Mounted Camera System Integrates Detailed Behavioral Monitoring with Multichannel Electrophysiology in Freely Moving Mice. Neuron, 100(1):46–60.e7, October 2018.

[21] Jérome Carriot, Mohsen Jamali, Maurice J. Chacron, and Kathleen E. Cullen. The statistics of the vestibular input experienced during natural self-motion differ between rodents and primates: Natural vestibular input in rodents and monkeys. The Journal of Physiology, 595(8):2751–2766, April 2017.

[22] Guy Bouvier, Yuta Senzai, and Massimo Scanziani. Head Movements Control the Activity of Primary Visual Cortex in a Luminance-Dependent Manner. Neuron, 108(3):500–511.e5, November 2020.

[23] Mateo V élez-Fort, Edward F. Bracey, Sepiedeh Keshavarzi, Charly V. Rousseau, Lee Cossell, Stephen C. Lenzi, Molly Strom, and Troy W. Margrie. A Circuit for Integration of Head- and Visual-Motion Signals in Layer 6 of Mouse Primary Visual Cortex. Neuron, 98(1):179–191.e6, April 2018.

[24] Ioana Medrea and Kathleen E. Cullen. Multisensory integration in early vestibular processing in mice: the encoding of passive vs. active motion. Journal of Neurophysiology, 110(12):2704–2717, December 2013.

